# The influence of bee movements on patterns of pollen transfer between plants: an exploratory model

**DOI:** 10.1101/2025.05.19.654816

**Authors:** Juliane Mailly, Thomas Besognet, Mathieu Lihoreau, Louise Riotte-Lambert

## Abstract

Most -if not all- pollinators make foraging decisions based on learning and memory. In interaction with environmental conditions and competitive pressure, pollinators’ cognition shapes their movement patterns, which in turn determine pollen transfers. However, models of animal-mediated pollination often make simplifying assumptions about pollinator movements, notably by not incorporating learning and memory. Better considering cognition as a driver of pollinators’ movements may thus provide a powerful mechanistic understanding of pollen dispersal. In this exploratory study, we connect pollinator behaviour and plant reproduction by using an agent-based model of bee movements implementing reinforcement learning. Simulations of two bees foraging together in environments containing twenty flowers shows how learning can improve foraging efficiency as well as plant pollination quality through larger mating distances and smaller self-pollination rates, while creating spatially heterogeneous pollen flows. This suggests that pollinators’ informed foraging decisions contribute to genetic differentiation between plant subpopulations. We believe this theoretical exploration will pave the way for a more systematic analysis of animal-mediated plant mating patterns, as model predictions can be tested experimentally in real bee-plant systems.

## Introduction

Accurately predicting pollination processes is a key challenge for sustainable food production and the conservation of natural ecosystems, as about 70% of flowering plants rely on pollination mediated by animals (Ollerton et al. 2011). Most - if not all - pollinating insects, birds and mammals rely on sensory cues, learning, and memory to visit nectar producing flowers, resulting in complex movement patterns (Fagan et al. 2013). One of the most striking manifestations of these sophisticated behaviours is trapline foraging, by which individual animals develop multi-destination routes to revisit known feeding locations in a stable order (bees: Janzen 1971; Thomson et al. 1997; Ohashi and Thomson 2009; Lihoreau et al. 2013; hummingbirds: Tello-Ramos et al. 2015; bats: Machado et al. 1998). Learning and memory also influence how animals react to competitive pressure, notably by enabling individuals (Riotte-Lambert et al. 2015) or groups (Aarts et al. 2021; Morinay et al. 2023) to segregate spatially and exploit different foraging areas. Such resource partitioning has been documented in different pollinator species (bees: Pasquaretta et al. 2019; bats: Goldshtein et al. 2020). In the end, the complex yet non-random movement patterns guided by pollinators’ cognition have profound effects on pollen dispersal, ultimately influencing plant mating patterns and fitness in a non-trivial way (e.g., Becher et al. 2016). Better understanding these cascading effects and being able to predict them in the long term is thus critical to better manage pollination services, for instance in protected areas or for precision agriculture, especially in the context of a looming pollination crisis (Goulson et al. 2015; Aizen et al. 2022).

Developing a mechanistic framework to connect pollinators’ movements to plant mating patterns has been identified as a priority for pollination ecology in the 21st century (Mayer et al. 2011). Spatially explicit individual-based modelling offers a powerful tool to understand, explore and predict this relationship (Mailly et al. 2025a). However, many models rely on oversimplifications of pollinator movements, assuming for instance that they perform correlated random walks or Lévy flights (e.g., Vallaeys et al. 2017), follow advection-diffusion processes (e.g., Capera-Aragones et al. 2021) or only use local perception (nearest-neighbour movements: e.g. Everaars et al. 2018, random walk guided by vision: e.g., Dorin et al. 2018). Some models have considered memory use (short-term memory to avoid revisits and flower constancy: e.g. Dorin et al. 2022; specific location stored in the long-term memory: MacQueen et al. 2022) and chemical communication (e.g., Dorin et al. 2018). However, to our knowledge, only Ohashi and Thomson (2009) considered reinforcement learning, whereby bees learn and modify their foraging decisions based on their past experience of nectar rewards. In this pioneering study, two behavioural strategies were tested: area-restricted search (stochastic movements, but travelling long distances after encountering low rewards) and sample-and-shift traplining (traplining, but sampling novel patches and shifting to neighbouring rewarding patches after encountering low reward). Their simulations showed that trapliners included into their foraging routes flowers that were more spread out compared to searchers, which increased plants’ mating distances. Trapliners also visited more flowers before returning to a given flower, which increased plants’ mate diversity and decreased self-pollination rates. Therefore, highest levels of pollination quality were reached in populations of pollinators with a high proportion of trapliners. While this is an important first step to understand the impact of cognitively guided movements on plant reproduction, this precursor work only compared two very different foraging strategies, and did not evaluate the impact of learning. It also did not investigate the impact of competition on pollen transport and plant reproduction.

Therefore, here we explored the predictions of a cognitively realistic, experimentally-validated, agent-based model of pollinators’ movements implementing learning and memory (Lihoreau et al. 2012; Reynolds et al. 2013; Dubois et al. 2021; Mailly et al. 2025b). In this model, foraging bees learn to exploit feeding locations based on reinforcement learning of flight vectors (Giurfa 2013; Chittka 2022), i.e., they learn by trial-and-error the relative values of transition movements linking pairs of feeding sites. This simple behavioural process can generate a wide range of foraging movements - from near random movements to traplining (Mailly et al. 2025b).With this mechanistic model, we explored how learning shapes the movement patterns of bees and the subsequent pollen dispersal by focusing on a simple scenario with two bees foraging in a meadow of twenty plants (but also providing a sensitivity analysis with more complex scenarios). As learning interacts with competition pressure in guiding bee movements (Makino and Sakai 2005; Ohashi et al. 2013; Pasquaretta et al. 2019), we also varied the nectar renewal duration of plants: Fast-renewing nectar should lessen the intensity of competition as plants are more likely to replenish in between visits, thus providing enough nectar for both bees, while slow-renewing nectar should prevent the bees from foraging on the same plants. For the sake of simplicity, in this exploratory study we did not vary other nectar traits such as nectar nutritional quality, caloric value or sugar concentration.

## Methods

### Bee movements

We used a model of bee movements as described in detail in (Mailly et al. 2025b). Briefly, our model simulates the behaviour of central-place foraging bees, which exploit a set of feeding sites (i.e. plants) from a stable nest location. At each step, bees decide to travel to a target location (a plant or the colony nest) based on positive and negative reinforcement learning processes underlying vector navigation in insects (Patel et al. 2024; Wolf 2024). This model and its variants have been previously used to simulate different scenarios with varying numbers of bees, plants and types of environments (Lihoreau et al. 2012; Reynolds et al. 2013; Dubois et al. 2021; Buatois et al. 2024; Mailly et al. 2025b).

#### Time

Time is represented by discrete 5-second timesteps.

#### Environment

The environment is a square area with continuous spatial coordinates. It contains the nest at (0; 0), and a set of plants. Each plant is characterised by its coordinates and the quantity of nectar it contains at each timestep. If the plant gets emptied by a bee, the nectar is renewed linearly in a fixed number of timesteps until reaching its maximal value. This maximal value is the same for all simulations (Table S1). The plant positions have x and y coordinates drawn from uniform distributions, which means that they are on average evenly spread out in the square environment. Importantly, as the dimension of the environment is a fixed parameter of the simulations, increasing the number of plants also increases the spatial density of plants. 25 randomly generated environment configurations were generated for each environmental condition (either 20 or 40 plants; see Simulation experiment paragraph below).

#### Behaviour

Bees move between plants or between the nest and a plant in a straight line, with constant speed. Their crop capacity is equivalent to the full nectar loads of 5 plants as in Dubois et al. 2021. Each bee performs a series of foraging bouts, which consist in leaving the nest and successively feeding on different plants until its nectar crop capacity is full or a maximum flying distance has been reached, and returning to the nest. The crop capacity and maximum distance are fixed parameters of the model (see Table S1). The bee then waits for a fixed amount of time in the nest before starting another foraging bout. We assume that the bee knows and remembers the position of all the plants in the environment and how to move between each pair of plants.

#### Learning and decision-making

When a bee makes a transition between plants A and B at time t, it perceives a value *V*_*t*_ (*A, B*) = *n*_*t*_ (*B*) × *p*_*d*_(*A, B*), where *n*_*t*_ (*B*) is the nectar available in plant B at time *t* and *p*_*d*_(*A, B*)approximates the probability of discovering plant B from plant A following a random walk. *p*_*d*_(*A, B*) is proportional to 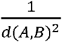, where *d*(*A, B*) is the distance between plants A and B, and it is normalised so that ∑ *p*_*d*_(*A*, .) equals 1 (see Dubois et al. 2021 for more information about this approximation). We thus assume that the initial difficulty to find plants modulates the subsequent value the bee allocates to transitions.

The bee’s expectation of a plant-to-plant transition value is initialised at 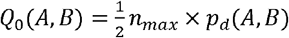, where *n*_*max*_ is the maximal nectar capacity of a plant. On subsequent realisations of the transition, the bee updates the expected value *Q*(*A, B*) so that *Q*_*t*_ (*A, B*) = *αV*_*t*_ (*A, B*) + (1 - *α*) *Q*_*t* -1_(*A, B*) where *α* is the learning rate of the bee, ranging between 0 and 1. If *α* is smaller than 0.5, the bee relies more on its previous expectation than on the newly experienced value. Conversely, if *α* is larger than 0.5, the bee relies more on the newly experienced value than on its previous expectation (Watkins and Dayan 1992).

Every time the bee visits a plant, it chooses the next plant to visit based on its expected values, using a softmax function defined as follows: When on plant A, the probability to choose to go to plant B is 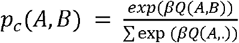. β ≥ 0 is the exploration-exploitation parameter. It controls how noisy a bee’s decision is. The closer to 0, the more the bee chooses ‘at random’ between the different options. The higher *β*, the more the bee tends to choose the transition with the largest expected nectar value.

#### Memory

We assume that bees have a long-term memory (i.e. memory lasting for the whole duration of the simulation) of the positions of the plants (Thomson 1996). This is realistic because we only simulate up to 40 different plants and both Thomson et al. (1997) and Makino and Sakai (2005) studied traplining behaviours by bees in environment configurations of 37 feeding sites. We also assume that they remember the expected values associated with each transition between plants. Although we are not aware of a direct experimental demonstration of such a hypothesis, bees are well known to associate individual food sources with reward qualities (Cnaani et al. 2006), develop long-term expectations of rewards (Gil et al. 2007), and they have been experimentally observed to rapidly abandon flight vectors leading to non-rewarding feeding sites (Lihoreau et al. 2012), making this assumption plausible. The bees also have a short-term (i.e. working) memory that inhibits the return to plants visited a short time ago (see Menzel 2009 for a review). The span of the working memory represents the time during which recently visited plants are excluded from the transition options in the bees’ decision-making process. This span was fixed at 60 seconds during the simulations. Spatial working memory abilities have been experimentally demonstrated in honey bees (Brown and Demas 1994; Brown et al. 1997) and bumble bees (Boisvert et al. 2007; Samuelson et al. 2016).

### Pollen dispersal

We added a pollen transfer module to the behavioural model. This pollination module relies on a geometric pollen carryover function (Montgomery 2009). Each time a bee visits a plant, it picks up 1 arbitrary unit of pollen and drops a proportion *p* of its pollen crop - called the pollen deposition rate - thereby depositing *p* pollen units from the previously visited plant, *p*(1 − *p*) pollen units from the second-to-last visited plant, *p*(1 − *p*)^2^ from the third-to-last, etc. Therefore, higher values of *p* lead to a faster decline in pollen deposition on plants visited later in the foraging bout. Pollen transfers below 0.01 pollen units were neglected. We assumed that all plants are bisexual and carry an undepletable amount of pollen. We also assumed that there is no limit to the amount of pollen the bee carries during a foraging bout and that the bee grooms in the nest, so that there are no pollen residues when starting a new foraging bout.

### Simulation experiments

We simulated two bees foraging in an environment with 20 plants. We varied the duration of nectar renewal between 200 and 1200 seconds and the learning rate **α** (either 0. or 0.5). We chose renewal duration values of a similar order of magnitude as the duration of a bee’s foraging bout. We ran sensitivity analyses on the number of bees (either 1, 2 or 5), the number of plants (either 20 or 40), the exploration-exploitation parameter *β* (either 10. or 20.) and the pollen deposition rate *p* (either 0.25 or 0.5) (Fig. S1 to S8). The number of bees and plants were chosen such that all environments could theoretically provide enough nectar for all the bees. All other parameter values were fixed across simulations (see Table S1 for parameter values and justifications). Overall, all parameter values were kept constant within each simulation. We used 25 different randomly-generated environments for which we tested each parameter combination 50 times, resulting in a total of 1250 simulations per parameter combination. Each simulation lasted 40,000 seconds.

### Data analyses

We computed various metrics from the model outputs to characterise bee foraging efficiency, movement patterns, and plant pollination quality.

## Metrics of foraging behaviour

1. Number of visits: The number of visits made by a bee during a foraging bout. This includes multiple visits to the same plant.
2. Nectar intake rate: The quantity of nectar per unit of time collected by a bee during a foraging bout in µL/sec.
3. Similarity index: This index quantifies to what degree two consecutive foraging bouts *a* and *b* share smaller plant visitation subsequences (Dubois et al. 2021). It computes as: 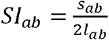 . Where *s*_*ab*_ is the total number of plant-to-plant transitions that compose the subsequences found in both *a* and *b*, and *l*_*ab*_ is the length of the longest sequence. The length of the subsequences was set at 3 (as in Dubois et al. 2021), which is long enough to detect subsequences but not too large compared to the length of a single foraging bout (5 full-plant visits, up to 13 plant visits on average). When two consecutive foraging bouts have no subsequence in common, the similarity index equals 0. If two consecutive foraging bouts are identical, the index equals 1.
4. Local intensity of competition: This index of space partitioning evaluates how much the plants exploited by an individual were also exploited by competing foragers (Riotte-Lambert et al. 2017). It is computed during a time window consisting of the last 5,000 seconds of the simulation. For an individual *i* and a time window *j*, the local intensity of competition *I*_*loc*_[*i*]_*j*_ is computed as 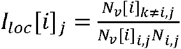 Where *N*_*i*,*j*_ is the number of plants visited at least once by individual *i* during the time window *j*, and *N*_*v*_ [*i*] _*i*,*j*,_ and *N*_*v*_ [*i*] _*k ≠ i*,*j*,_ are the total number of times any of the *N*_*i*,*j*,_ plants visited by *i* and by any individual other than *i* respectively during the time window *j*.
5. Spatial distribution of foragers: We computed the average number of visitors per plant during the last 5,000 seconds of the simulation. The data from all environmental configurations were spatially interpolated by using a nearest-neighbour interpolation (SciPy’s interpolate sub-package in Python; Virtanen et al. 2020). This allowed us to go from a discrete spatial representation of data (each datapoint is associated to a discrete plant position) to a continuous spatial representation that can be plotted in a rasterised 2D heatmap for visualisation.

The local intensity of competition and the spatial distribution of foragers were computed for the last 5,000 seconds of the simulation. The other metrics (number of visits, nectar intake rate and similarity index) were computed for the 29th foraging bout as the subsequent bouts were not present in all simulations. We focused on the end of the simulation so that bees were given enough time to learn and stabilise routes. Metrics were then averaged over all bees (except for the spatial distribution of foragers) and simulations for each parameter combination, except as noted.

### Metrics of pollination patterns

All metrics were computed for the last 5,000 seconds of the simulation and averaged over all plants (except for population-wide metrics) and simulations for each parameter combination, except as noted.

1. Mating distance: The median mating distance of the whole plant population, weighted by the amount of pollen transferred (Ohashi and Thomson 2009).
2. Self-pollination rate: The ratio between the amount of a plant’s own pollen that it received to the total amount of pollen received (Ohashi and Thomson 2009).
3. Mate diversity: The number of donor plants per recipient plant (adapted from Ohashi and Thomson 2009).
4. Spatial segregation of pollen dispersal: We computed the modularity *M* of the weighted bipartite network between donor plants and recipient plants - a method widely applied to plant-pollinator networks (e.g., Brimacombe et al. 2022; Gay et al. 2024). This undirected network is defined with two sets of nodes: the set of donor plants (i.e., all the plants that have successfully sent pollen to at least one plant) and the set of recipient plants (i.e., all the plants that have received pollen from at least one plant). These two sets of plants can overlap but are decomposed in this network. The weight of the edge between a donor and a recipient plant is the pollen quantity transferred from the donor to the recipient. We then computed the modularity *M* of this network, which describes how clustered the network is. This index was computed with the *DIRTLPAwb+* algorithm (Beckett 2016) using the R package “bipartite”. To allow for comparisons between networks, we normalised *M* so that it ranged between 0 and 1 (see details of this normalisation in Pasquaretta and Jeanson 2018). If *M* equals 1, then each donor plant sends its pollen to an isolated set of recipient plants. If *M* equals 0, all donor plants send their pollen to the same set of recipient plants.

## Results

### Effect of learning and nectar renewal duration on bee foraging

Foragers visited more plants as the nectar renewal duration increased, but this effect was much stronger for non-learning than for learning bees (Fig. 1.A). Overall, learning bees visited fewer plants per foraging bout (Fig. 1.A). Nectar intake rates decreased when nectar renewal duration increased, but less steeply for learning than for non-learning bees (Fig. 1.B). Learning bees had a higher average intake rate than non-learning bees. The similarity index, which measures the repeatedness in plant visitation sequences, was higher for learning bees than for non-learning bees, and decreased with increasing nectar renewal durations (Fig. 1.C). By contrast, the similarity index for non-learning bees remained constant for all values of nectar renewal durations (Fig. 1.C). Finally, learning bees increased their spatial segregation as nectar renewal duration increased, as shown by the decrease in local intensity of competition values (Fig. 1.D). However, the local intensities of competition experienced by non-learning bees remained high for all values of nectar renewal durations (Fig. 1.D).

**Figure 1.**
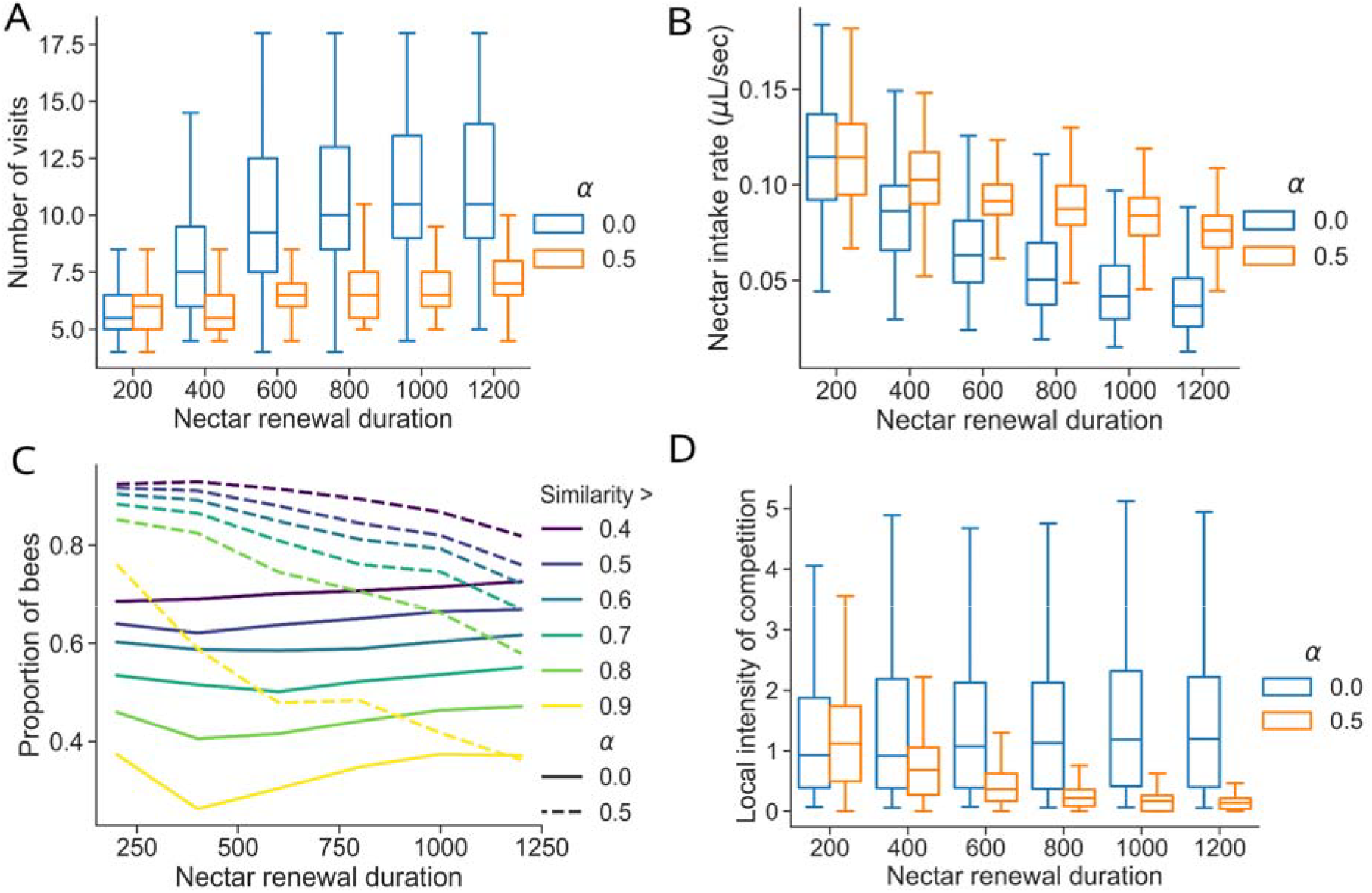
Effect of learning rate (colour in panels A-B-D, line style in panel C) and nectar renewal duration (x-axis) on metrics of bee foraging behaviour (y-axis): (A) number of plant visits per foraging bout, (B) nectar intake rate of a bee (C) proportion of bees across all simulations whose bout similarity index value was above a certain threshold (by colour), (D) local intensity of competition. In (A-B-D), the variability is shown across the 1250 simulations (50 replicates on 25 different environment configurations). More details about simulations and metric computation in the methods.

The spatial segregation of bees was also apparent when looking at the number of foragers visiting each plant. The learning condition led to a smaller average number of visitors per plant than the non-learning condition (Fig 2.A and C), which means that the two competing bees tend to forage on different plants. This trend increased with nectar renewal duration (Fig. 2.A). Non-learning bees both visited all plants on a broad area around the nest, and this pattern was not affected by nectar renewal duration (Fig. 2C).

**Figure 2.**
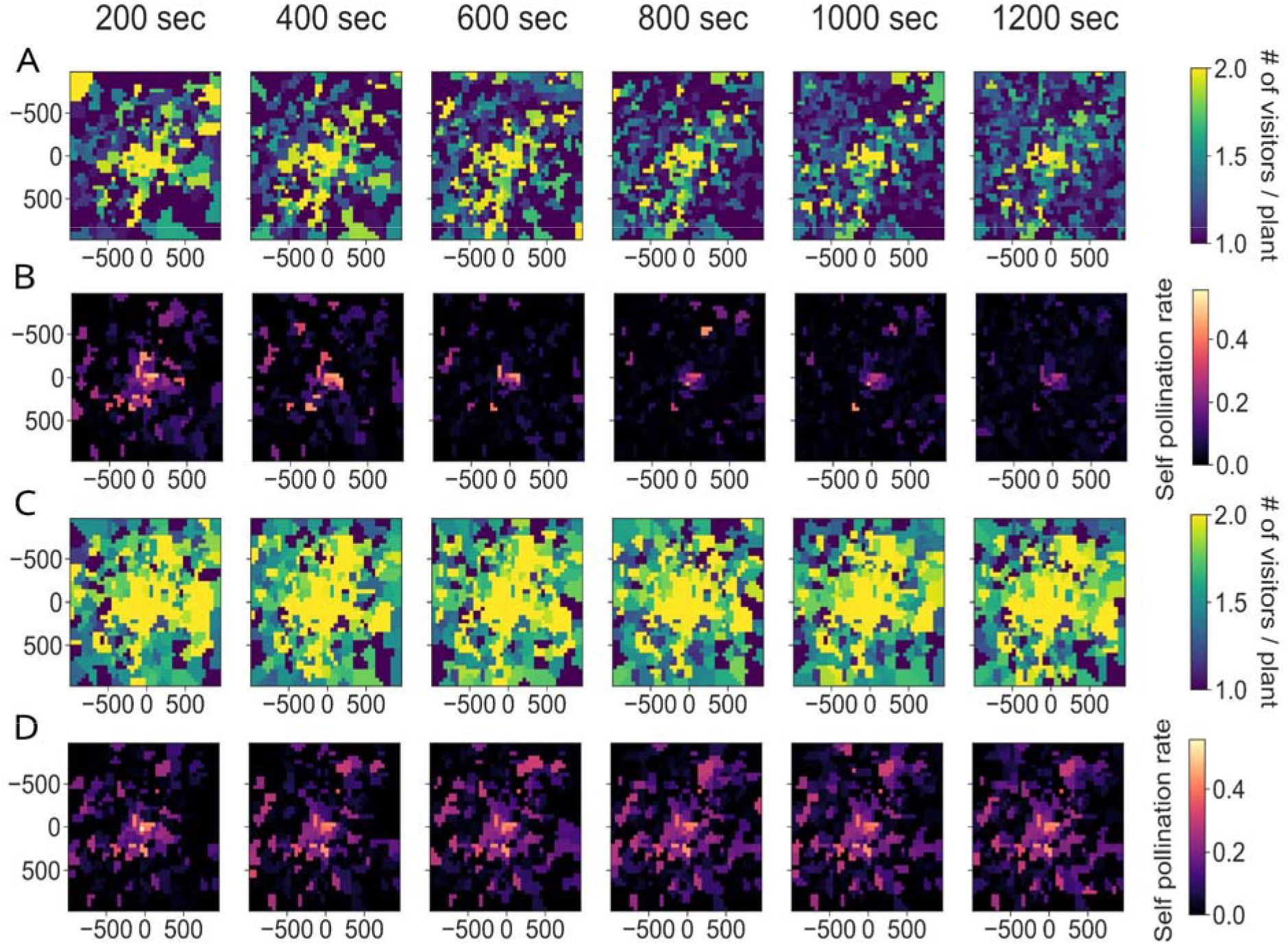
Effect of learning rate (for panels (A-B) and for panels (C-D)) and nectar renewal duration (by column) on (A-C) the spatial distribution of the average number of visitors per plant across simulations and (B-D) the spatial distribution of the average self-pollination rates per plant across simulations. The heatmap is based on the spatial interpolation of all plants from the 25 different environment configurations, averaged across 50 different simulations. See details about simulations and metric computation in the methods.

### Effect of learning and nectar renewal duration on plant pollination

The mating distances of plants pollinated by learning bees increased with nectar renewal duration (Fig. 3A). By contrast, the mating distances of plants pollinated by non-learning bees remained consistently lower for all nectar renewal duration values (Fig. 3.A). The self-pollination rate decreased with nectar renewal durations in the presence of learning bees (Fig. 3.B). Only plants close to the nest displayed a high self-pollination rate (Fig. 2.B). By contrast, in the presence of non-learning bees, the self-pollination rate stayed high for all nectar renewal durations and for most plants in the environment (Fig. 3.B, Fig. 2.D). Mate diversity increased with nectar renewal durations whether the bees were learning or not, but values were on average higher for plants pollinated by non-learning bees (Fig. 3.C). Finally, in the presence of learning bees, the donor-recipient plant network modularity increased with increasing nectar renewal durations, but decreased in the presence of non-learning bees (Fig. 3.D). This increase in modularity with learning bees likely derived from the increase in spatial segregation (Fig. 1D). By contrast, the decrease in modularity with non-learning bees likely derived from the conjunct effect of the steep increase in the number of plants visited (Fig. 1A) and the absence of spatial segregation (Fig. 1D).

**Figure 3.**
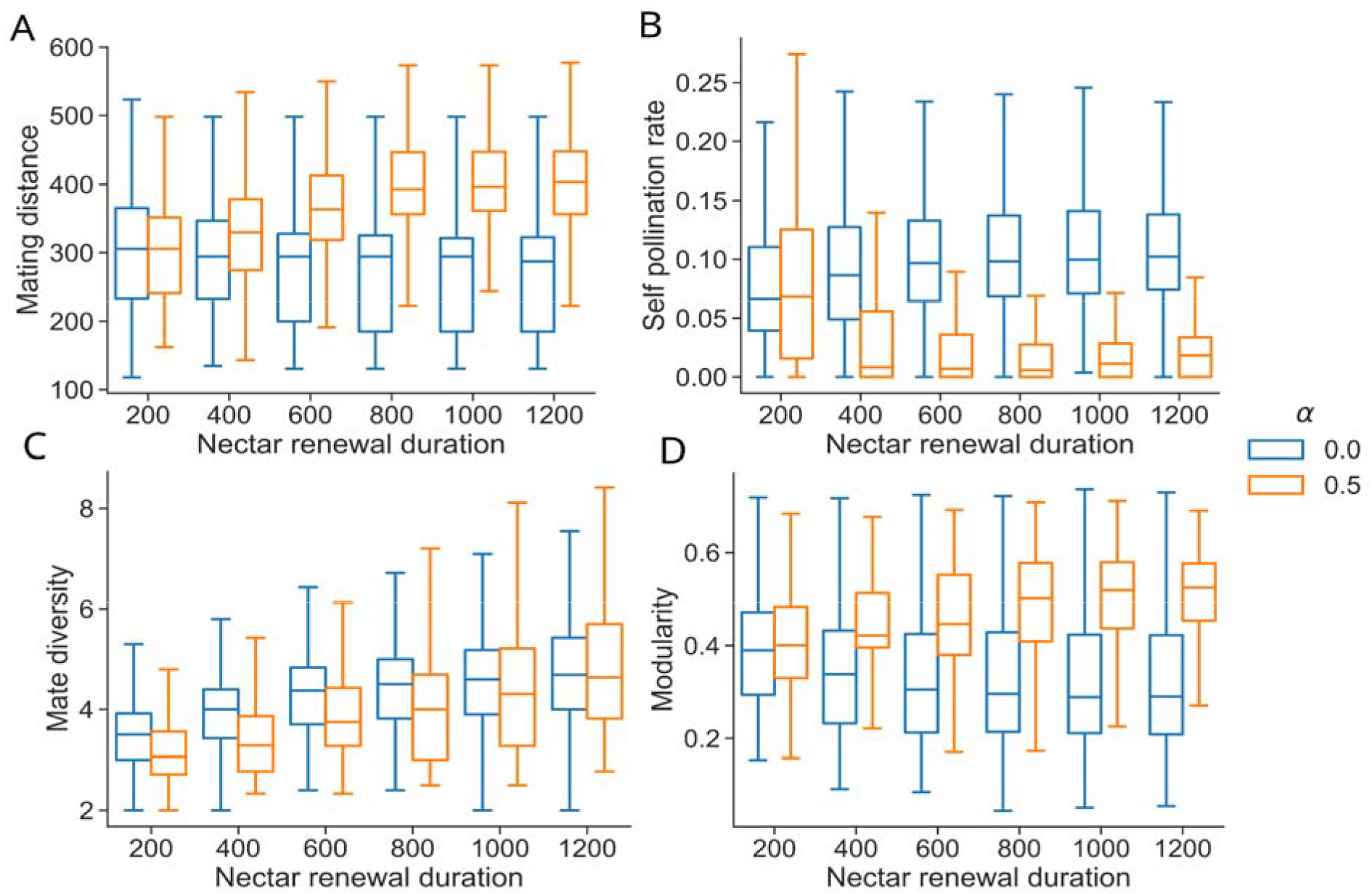
Effect of learning rate (by colour) and nectar renewal duration (x-axis) on metrics of plant pollination patterns (y-axis): (A) median plant population mating distance, (B) self-pollination rate of a plant, (C) mate diversity of a plant, (D) modularity of the donor-recipient plant bipartite network. The variability is shown across the 1250 simulations (50 replicates on 25 different environment configurations). More details about simulations and metric computation in the methods.

### Sensitivity analyses

The following parameters did not qualitatively impact the previously described effects of learning and nectar renewa duration, except as noted.

#### Effect of the number of bees (1 vs. 2 vs. 4 bees)

Whether or not bees learn, when the number of bees increases each bee tends to visit more plants per bout (Fig. S1) and the average individual nectar intake rate decreases (Fig. S2). The number of bees did not impact the average similarity index when bees were not learning (Fig. S3). However, when bees were learning, the similarity index decreased when the number of bees increased (Fig S3). In the 4-bees condition, the similarity index of learning bees was on average lower than that of non-learning bees for long renewal duration values (Fig. S3). A higher number of bees increased the local intensity of competition (Fig. S4).

The number of bees had a negligible effect on plant mating distances (Fig S5). Increasing the number of bees increased the self-pollination rate of plants pollinated by learning bees but not of plants pollinated by non-learning bees (Fig S6). Increasing the number of bees led to an increase of mate diversity in both conditions (Fig. S7). In the 4-bees condition with 20 plants, the difference in mate diversity between learning and non-learning conditions is more pronounced with higher mate diversity values in the presence of learning bees (Fig S7).

Increasing the number of bees led to a slight decrease of modularity in the non-learning condition (Fig. S8). When bees were learning and the pollen deposition rate was low, increasing the number of bees increased the modularity but with a high pollen deposition rate, increasing the number of bees led to a slight decrease in modularity. In the 1-bee condition, the learning and non-learning conditions led to equivalent levels of modularity, with sometimes non-learning bees leading to slightly higher levels of modularity (when and 40 plants). Interestingly, when there were only 20 plants for 4 learning bees (i.e., the environment was saturated), the modularity decreased slightly with increasing nectar renewal durations. In the presence of 4 non-learning bees, the modularity levels stayed rather equivalent for all nectar renewal durations (Fig S8).

#### Effect of the number of plants (20 vs. 40 plants)

Increasing the number of plants led to higher nectar intake rates (Fig S2), smaller similarity index values (Fig S3), and smaller local intensity of competition values - i.e. more spatial segregation between foragers (Fig S4). For plants, increasing the number of plants led to shorter plant mating distances (Fig. S5), smaller self-pollination rates (Fig. S6) and higher modularity values (Fig S8). Overall, the effect of learning on the local intensity of competition (Fig. S4), and on pollination metrics was more pronounced for the 20-plants condition.

#### Effect of the exploration-exploitation parameter *β* (10 vs. 20)

Increasing the value of *β* led to an increase in similarity index values (Fig S3) and in the local intensity of competition (Fig S4). Increasing *β* also led to more visible differences in modularity values between the learning and non-learning conditions (Fig S8).

#### Effect of the pollen deposition rate p (0.25 vs. 0.5)

At the plant level, higher pollen deposition rates favoured pollen transfer between two successively visited plants within a given foraging bout, with fast-decreasing transferred pollen amounts to subsequently visited plants (see Methods section). Therefore, a higher pollen deposition rate led to smaller mating distances (Fig S5) and self-pollination rates (Fig S6). However, a higher pollen deposition rate did not affect mate diversity (Fig S7) - likely because our measure of mate diversity did not account for the quantity of pollen transferred. Finally, a higher pollen deposition rate led to higher modularity values (Fig S8), which means that the spatial network of pollen dispersal is structured in more pollen clusters.

## Discussion

Pollination models often rely on simplifying assumptions on pollinators’ movements and their use of sensory cues, learning and memory (Mailly et al. 2025a). Here, building on previous work (e.g., Ohashi and Thomson 2009; Dorin et al. 2018; Kortsch et al. 2023), we introduced a model of pollen dispersal implementing movements based on more realistic cognitive abilities (Mailly et al. 2025b). At the pollinator level, we confirm that reinforcement learning enables individuals to increase their foraging efficiency by partly escaping competition through spatial segregation (Dubois et al. 2021). At the plant level, pollination by foragers with learning abilities improves average pollination quality with increased plant mating distances and lower rates of self-pollination.

Interestingly, our simulations show how simple learning rules that have been well-described in many pollinator species can simultaneously benefit both pollinators and plants. Without learning, bees’ food intake rate steeply declines when nectar renewal duration increases (Fig. 1.B). However, learning bees can much better cope with high nectar renewal durations (Fig. 1.B), reaching high levels of foraging efficiency in part through spatial segregation (Fig. 1.C). In our simulations, learning is also extremely beneficial for the plants as learning-bee pollination leads to higher mating distances and lower self-pollination rates (Fig. 3.A and B), and rather equivalent mate diversity (Fig. 3.C), compared to non-learning-bee pollination. These effects of bee learning on plant reproduction partly align with the predictions of Ohashi and Thomson (2009) that plants pollinated by trapliners (as opposed to random foragers) tend to have larger mating distances and mate diversities and smaller self-pollination rates. However, our results also suggest the new possibility that plant nectar renewal rates might be subjected to a trade-off. Plants attractivity is likely to be inversely related to nectar renewal duration, as pollinators’ intake rates decrease when nectar renews more slowly (Fig. 1.B). On the other hand, nectar production is costly (Pyke and Ren 2023), and, in the presence of learning bees, slower nectar renewal leads to larger mating distances, higher mate diversity and smaller self-pollination rates (Fig. 3.A, 2.B and 2.C), while leading to a potentially lower total nectar production over the plant’s lifetime. Recent evidence suggests that this floral trait is highly evolvable, and thus has the potential to respond to such pollinator-mediated selection (Romero-Bravo and Castellanos 2024).

Our model also suggests the intriguing possibility that pollinator learning can lead to subnetworks of pollen flow by which pollen is only shared between some individuals within the plant population (Fig. 3.D). In the long term, such a clustering of pollen flows could favour genetic differentiation in the plant population (Ohsawa et al. 1993; Linhart and Grant 1996; Wessinger 2021). However, this effect should be verified over several plant generations, as isolated subpopulations might cross again in the next season. High pollen deposition rates might also lead to more genetically isolated plant subpopulations, as it decreases mating distance and increases pollen flow clustering (Fig. S5 and S8). To avoid inbreeding, plants could develop strategies to decrease pollen deposition rates by promoting pollen adhesion to pollinating animals (e.g., a coating called pollenkitt; Lin et al. 2013).

Although our model is based on behavioural data and has been shown to successfully replicate bees’ plant visitation sequences in various experimental conditions (Lihoreau et al. 2012; Reynolds et al. 2013; Dubois et al. 2021), it remains exploratory at this stage. Its predictions serve as qualitative investigative directions and should be tested and validated experimentally. In particular, our approach presents several simplifying hypotheses that were necessary for such an exploratory work. The simulated environment is an abstraction from real landscapes, only considering the position of plants or feeding sites and neglecting landmarks that might shape movements (Collett and Collett 2002; Brebner et al. 2021). We also chose a simple geometric pollen carryover function, which only depends on one parameter (p) and is thus easy to compute and parametrise. However, experimental work suggests that this function underestimates the amount of pollen deposited after several visits (Morris et al. 1994). Our results thus likely underestimate the number of connections in the plant population network, which could result in underestimations of plants’ mating distance, mate diversity and overestimations of the modularity of the network. We also assume an unlimited pollen production in plants and no limitation in pollen transport by bees. In reality, bees deplete the pollen of a plant after only a couple of visits (Harder and Thomson 1989) and pollen grains compete for space on pollinators (Moir and Anderson 2023; Santana et al. 2025), which could lead to shorter pollen flow distances. A way forward would thus be to implement both pollen depletion on plants and pollen transport limitation on bees (Harder and Wilson 1998). Our model is also an abstraction of bee cognition that overlooks mechanistic navigational mechanisms such as spatial memory building, path integration or visual guidance that can be sources of direction biases or navigational errors in real foragers (e.g., Le Moël et al. 2019).

The simplicity of the decision rules implemented in our model would make it highly adaptable to new research questions with minimal changes. For example, future work could explore the impact of plant spatial aggregation (Cresswell 1997; Comba 1999; Osborne and Williams 2001). Habitat patchiness can be considered at several levels: between plants, by aggregating them into patches, or within plants, by representing several flowers belonging to a common plant. Moreover, other floral traits could be added into the model. For example, nectar nutritional value could be incorporated into a nutritional geometric framework (e.g., Helm et al. 2017) to model pollinators’ movements, where pollinators would seek to optimise their own diet or the colony’s nutritional state (Lihoreau et al. 2017). In addition, different forms of communication could be implemented, whether direct like the waggle dance (e.g., Dornhaus et al. 2006) or indirect like scent marking on plants (e.g., Dorin et al. 2018). These different forms of communication would greatly impact how foragers respond to competition but also their individual learning experiences, as they could exploit more easily profitable feeding sites (with the waggle dance) or avoid depleted plants (using scent marking). Through these adjustments, the model could represent different bee species (e.g. Buatois et al. 2024). Territorial bird pollinators could also be modelled by implementing aggressive behaviour between individuals (Krauss et al. 2017). In general, many species could be captured by the model by changing parameters such as the total foraging distance or the initial probabilities of transition between plants. These parameters would impact the foragers’ mobility, which is a key parameter that could influence pollen transfers (Wessinger 2021). Ultimately, this approach would enable the comparison of the effects of these different pollinator species on plant mating patterns and assess their functional complementarities (Garibaldi et al. 2013; Rader et al. 2016).

Ultimately, we believe this mechanistic approach has strong potential to advance fundamental knowledge in pollination ecology, for instance by improving predictions of long-term plant-pollinator network dynamics (Mailly et al. 2025a). If widely adopted, such models could also inform practical actions in conservation, helping to protect pollination services, or in precision agriculture, enhancing crop yield and limiting unwanted gene flow. Like all biological interactions, plant-pollinator relationships are complex (Peralta et al. 2024). A deeper understanding of local interactions–specifically, pollinator movements that mediate pollen dispersal–can reveal cascading effects across system levels and offer valuable insights into emerging topics such as pathogen spillover, pollination syndromes, and community resilience (Knight et al. 2018).

## Acknowledgments

This work was funded by the CNRS and the ERC Cog Bee-Move (grant number 101002644).

