## Supplementary material for "The influence of bee movements on patterns of pollen transfer between plants: an exploratory model": Fig. S

### Supplementary data

Table S1. Parameters used in each simulation and justifications.

| **Type of parameter** | **Parameter name** | **Description** | **Value** | **Justification** |
| --- | --- | --- | --- | --- |
| Simulations | Number of simulations | Number of independent simulations ran for each combination of parameter set and environment configuration. | 50 | Chosen to get 1250 simulations per parameter set (50 simulations in 20 different randomly-generated environment configurations) |
| Simulations | Number of seconds per timestep | Number of seconds represented by one simulation timestep. | 5 | Small enough temporal resolution to simulate bee movements between flowers. |
| Simulations | Duration of simulation | Duration of one simulation in seconds. | 40000 (s) | Time required for the stabilisation of bee behaviour |
| Environment generation | Number of flowers | Number of flowers in the environment. | 20 or 40 | Each flower represents 20% of the bee’s crop capacity (see “capacity of flowers” and “maximal crop capacity”). The number of flowers was chosen to ensure that the decisions of the bee are not constrained by a too low number of available flowers. |
| Environment generation | Foraging range | The patches of flowers are distributed in the environment such that their x,y coordinates vary within [-foraging range; foraging range]. The nest is always at (0,0). | 1000 (m) | Coherent with the foraging range of *Bombus terrestris* (Knight et al., 2005 [1]) |
| Environment generation | Number of environment configurations | Number of different environments randomly generated for each parameter set. The environments are generated as explained in the Methods. | 25 | Chosen to get 1250 simulations per parameter set (50 simulations in 20 different randomly-generated environment configurations) |
| Flowers | Capacity of flowers | Maximal nectar capacity of the flower in µL. | 10 (µL) | Corresponds either to a small patch of flowers or to an inflorescence (Luo et al., 2014 [2]; Adgaba et al., 2017 [3]) |
| Flowers | Duration of nectar replenishment | Duration (in seconds) a flower takes to get completely replenished in nectar, assuming a linear replenishment. After this delay, the flower will stay full until it gets visited by a bee. | Varied between 200 and 1200 sec | We chose values in coherence with the time spent in the nest and the working memory. In real life, flower renewal durations can vary between half an hour (e.g. Gilbert et al., 2001) [4] and several hours (e.g. Luo et al., 2014) [2] |
| Bee foraging | Number of bees | Number of bees foraging simultaneously. | Varied between 1 and 4 | Varied for the sensitivity analysis. |
| Bee foraging | Maximum distance travelled | Maximum distance a bee can fly before having to return to the nest. We assume here that the bee flies in a straight line between flowers. | 3000 (m) | Extracted from the raw data of Woodgate et al. (2016) |
| Bee foraging | Maximal crop capacity | Maximum amount of nectar in µL a bee can carry. | 50 (µL) | Honey bee crops contain around 50µL of nectar (e.g. Seeley, 1994 [5]); bumble bee crops contain around 100µL of nectar (e.g. Lihoreau et al., 2012 [6]). 50 corresponds to the nectar load of 5 full flowers (see ‘capacity of flowers’), which matches the experimental conditions of Lihoreau et al. (2012) [6]. |
| Bee foraging | Flight velocity | Flight velocity of the bee in m/s. | 5 (m/s) | From Riley et al. (1996) [7] |
| Bee foraging | Duration of nest stop | Time that the bee spends unloading nectar in the nest, in seconds. | 440 (s) | Extracted from the raw data of Woodgate et al. (2016) [8] |
| Bee foraging | Nectar foraging rate | Rate at which the bee absorbs the nectar in a flower, in µL/s. | 1 (µL/s) | From Pattrick et al. (2020) [9] |
| Bee cognition | Working memory span | Duration of the working memory in seconds, i.e. how long a flower is avoided by the bee after being visited. | 60 sec | Chosen as a very small value but sufficient to avoid endless loops of flower visits. |
| Bee cognition | α | Learning rate of the bee, i.e. the weight given to the newly observed value of the transition in comparison to the old expectation of that transition. It ranges between 0 (the bee doesn't learn after performing a transition), and 1 (the bee completely changes its expectation of the transition value to the most recent observation). | 0. or 0.5 | Chosen to compare learning and non-learning bees. For learning bees, $\alpha$equals 0.5 to match Riotte-Lambert et al. (2017) [10], i.e., a bee uses equally its past expectations and the new observation to update the expected value of a transition. |
| Bee cognition | β | Exploration-exploitation parameter, i.e. how much the bee relies on its own knowledge to make its decisions. It only takes positive values. The closer to 0, the more random the decisions of the bee. The larger the value, the more the bee follows its knowledge of the transition value and the less noisy the decision is. | 10 or 20 | Varied for the sensitivity analysis. |
| References | [1] Knight, M. E., Martin, A. P., Bishop, S., Osborne, J. L., Hale, R. J., Sanderson, R. A., & Goulson, D. (2005). An interspecific comparison of foraging range and nest density of four bumblebee (Bombus) species*. Molecular ecology*, 14(6), 1811-1820.  [2] Luo, E. Y., Ogilvie, J. E., & Thomson, J. D. (2014). Stimulation of flower nectar replenishment by removal: A survey of eleven animal-pollinated plant species. *Journal of Pollination Ecology*, 12, 52–62.  [3] Adgaba, N., Al-Ghamdi, A., Tadesse, Y., Getachew, A., Awad, A. M., Ansari, M. J., Owayss, A. A., Mohammed, S. E., & Alqarni, A. S. (2017). Nectar secretion dynamics and honey production potentials of some major honey plants in Saudi Arabia. *Saudi journal of biological sciences*, 24(1), 180–191.  [4] Gilbert, F., Azmeh, S., Barnard, C., Behnke, J., Collins, S. A., Hurst, J., ... & Course, T. B. E. F. (2001). Individually recognizable scent marks on flowers made by a solitary bee. *Animal behaviour*, 61(1), 217-229.  [5] Seeley, T. D. (1994). Honey bee foragers as sensory units of their colonies. *Behavioral Ecology and Sociobiology*, 34, 51-62.  [6] Lihoreau, M., Raine, N. E., Reynolds, A. M., Stelzer, R. J., Lim, K. S., Smith, A. D., . . . Chittka, L. (2012). Radar Tracking and Motion-Sensitive Cameras on Flowers Reveal the Development of Pollinator Multi-Destination Routes over Large Spatial Scales. *PLoS Biology,* 10, 1-13.  [7] Riley, J., Smith, A.D., Reynolds, D., Edwards, A.S., Osborne, J., Williams, I., Carreck, N. & Poppy, G. (1996). Tracking bees with harmonic radar. *Nature*. 379. 29-30.  [8] Woodgate, J. L., Makinson, J. C., Lim, K. S., Reynolds, A. M., & Chittka, L. (2016). Life-long radar tracking of bumblebees. *PLoS one*, 11, e0160333.  [9] Pattrick, J., Symington, H., Federle, W., & Glover, B. (2020). The mechanics of nectar offloading in the bumblebee *Bombus terrestris* and implications for optimal concentrations during nectar foraging. *Journal of The Royal Society Interface*. 17.  [10] Riotte-Lambert, L., Benhamou, S., Bonenfant, C., & Chamaillé-Jammes, S. (2017). Spatial memory shapes density dependence in population dynamics. *Proceedings of the Royal Society B: Biological Sciences*, *284*(1867), 20171411. | | | |

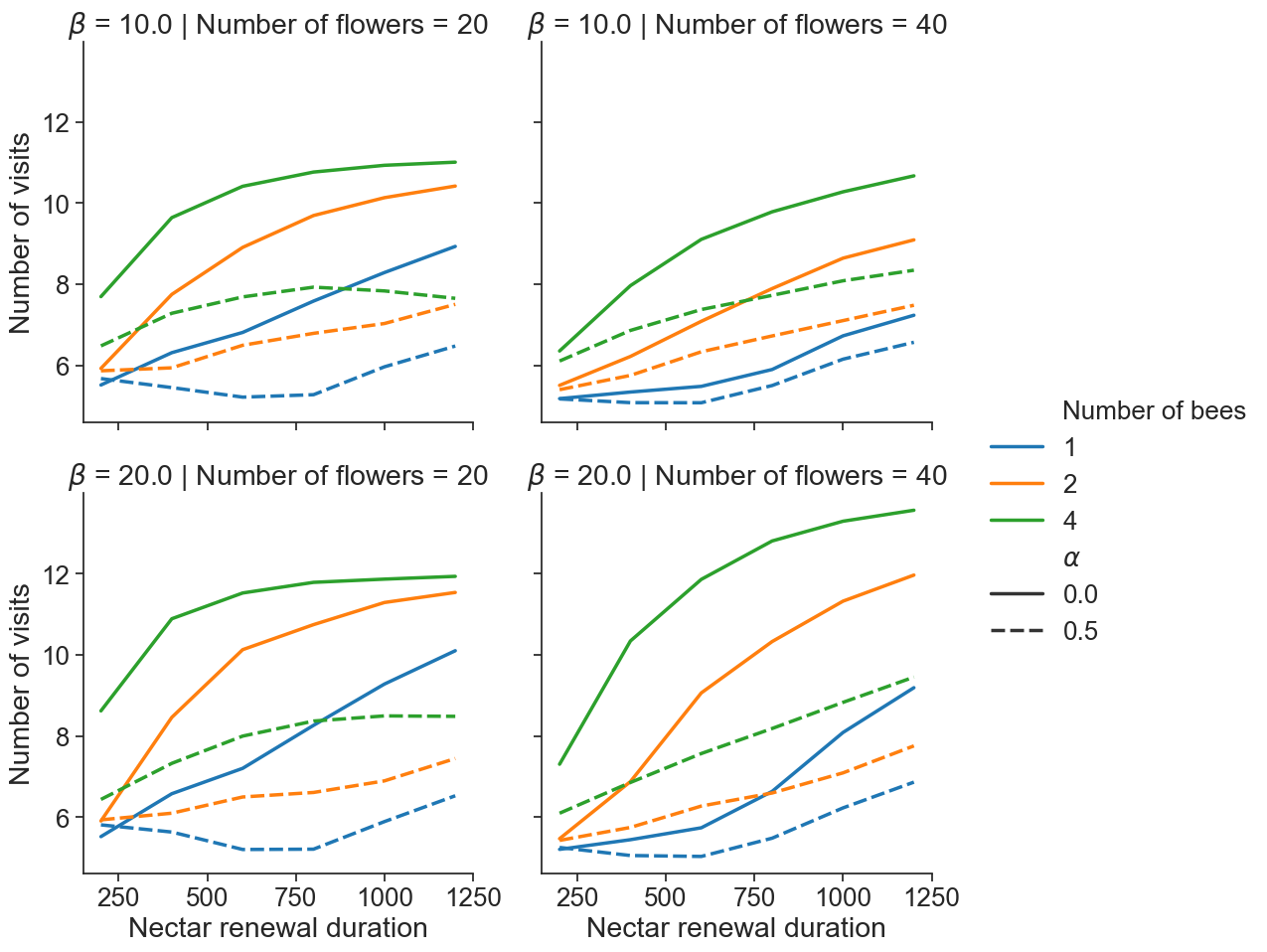

*Figure S1. Effect of learning rate* $\alpha$ *(line style), exploration-exploitation parameter* $\beta$ *(row), number of flowers (column), number of bees (line colour) and nectar renewal duration (x-axis) on average number of flower visits per foraging bout (y-axis).*

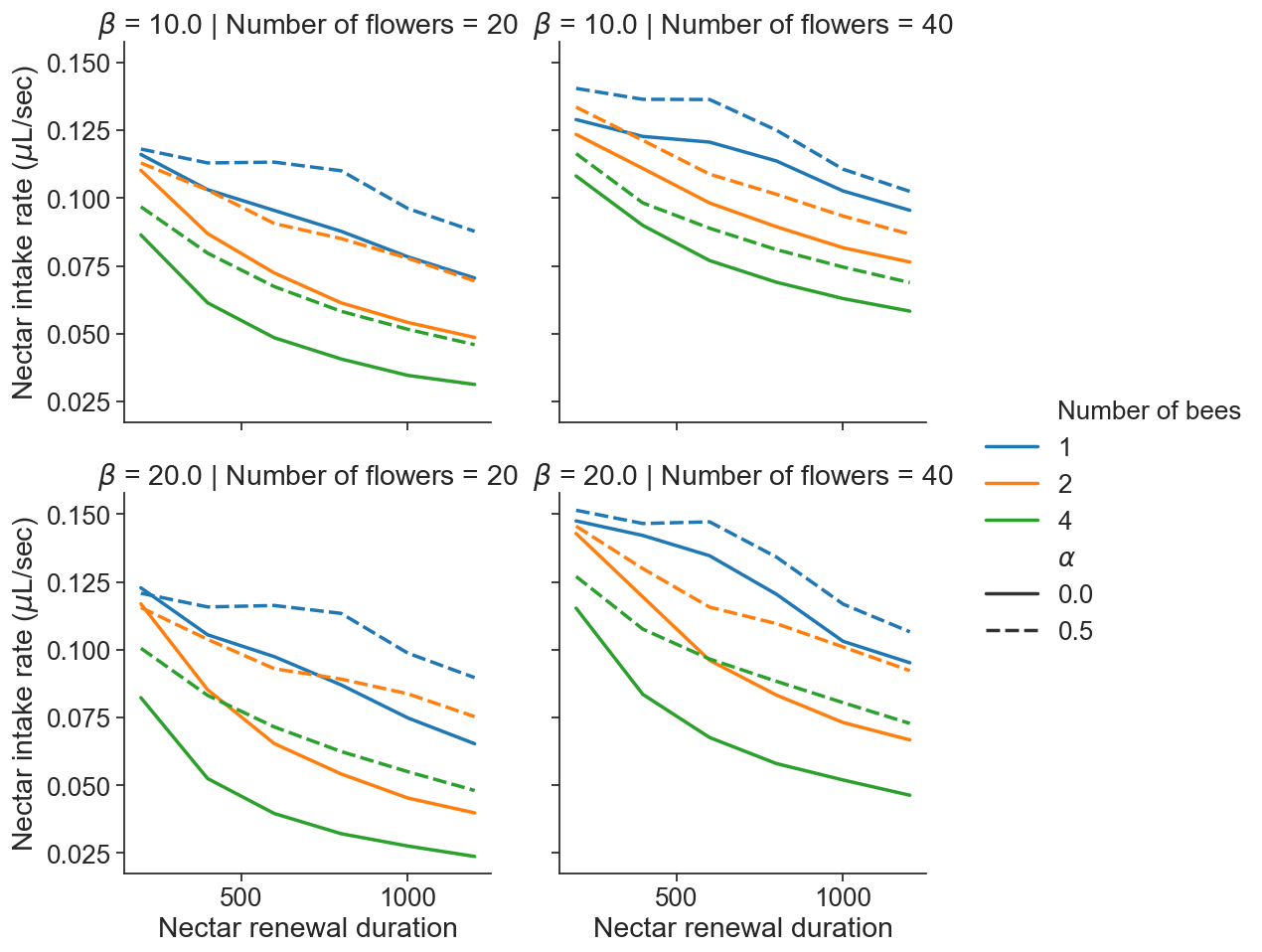

*Figure S2. Effect of learning rate* $\alpha$ *(line style), exploration-exploitation parameter* $\beta$ *(row), number of flowers (column), number of bees (line colour) and nectar renewal duration (x-axis) on average nectar intake rate of a bee (y-axis).*

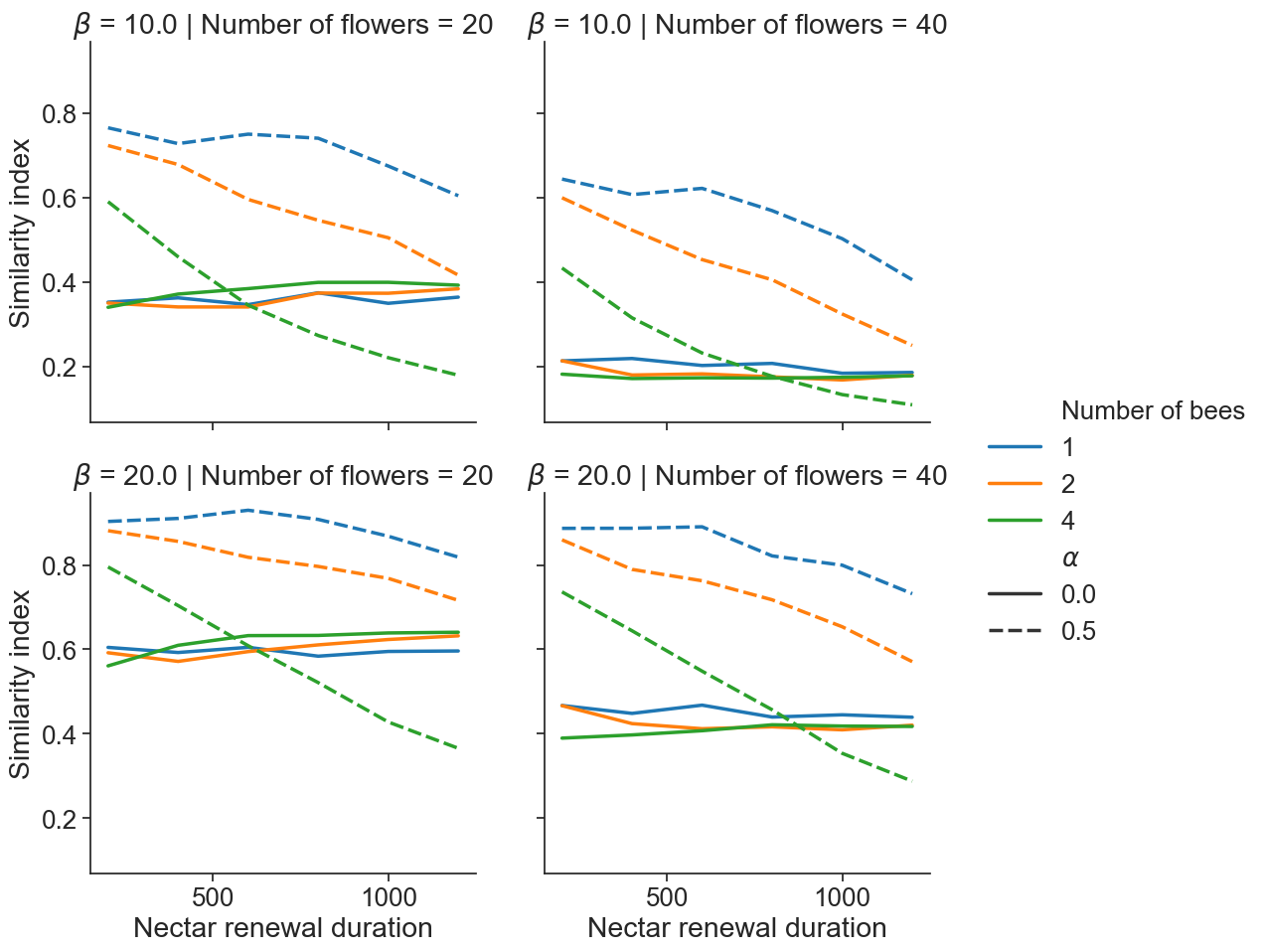

*Figure S3. Effect of learning rate* $\alpha$ *(line style), exploration-exploitation parameter* $\beta$ *(row), number of flowers (column), number of bees (line colour) and nectar renewal duration (x-axis) on average similarity index of a bee (y-axis).*

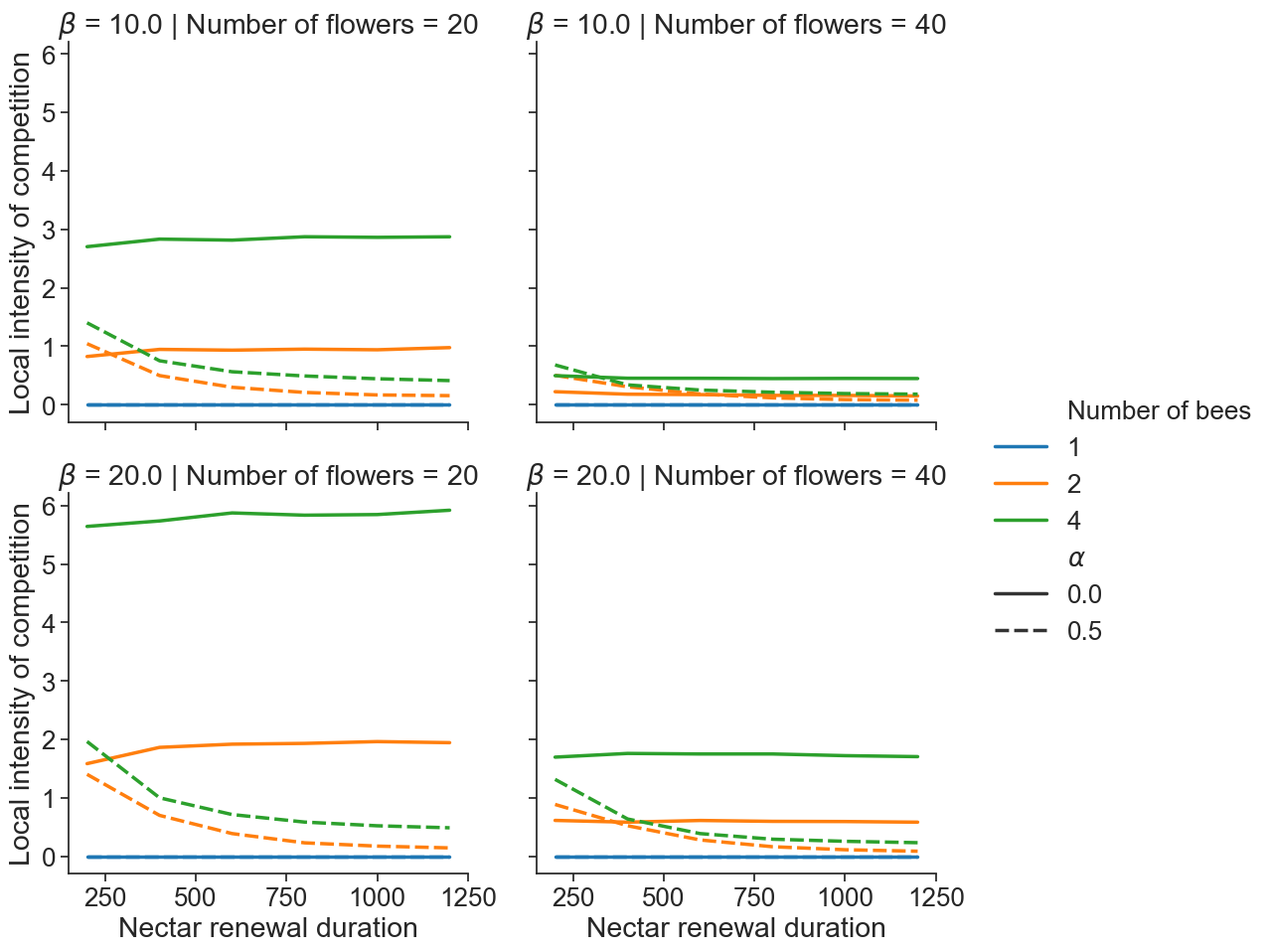

*Figure S4. Effect of learning rate* $\alpha$ *(line style), exploration-exploitation parameter* $\beta$ *(row), number of flowers (column), number of bees (line colour) and nectar renewal duration (x-axis) on average intensity of local competition of a bee (y-axis).*

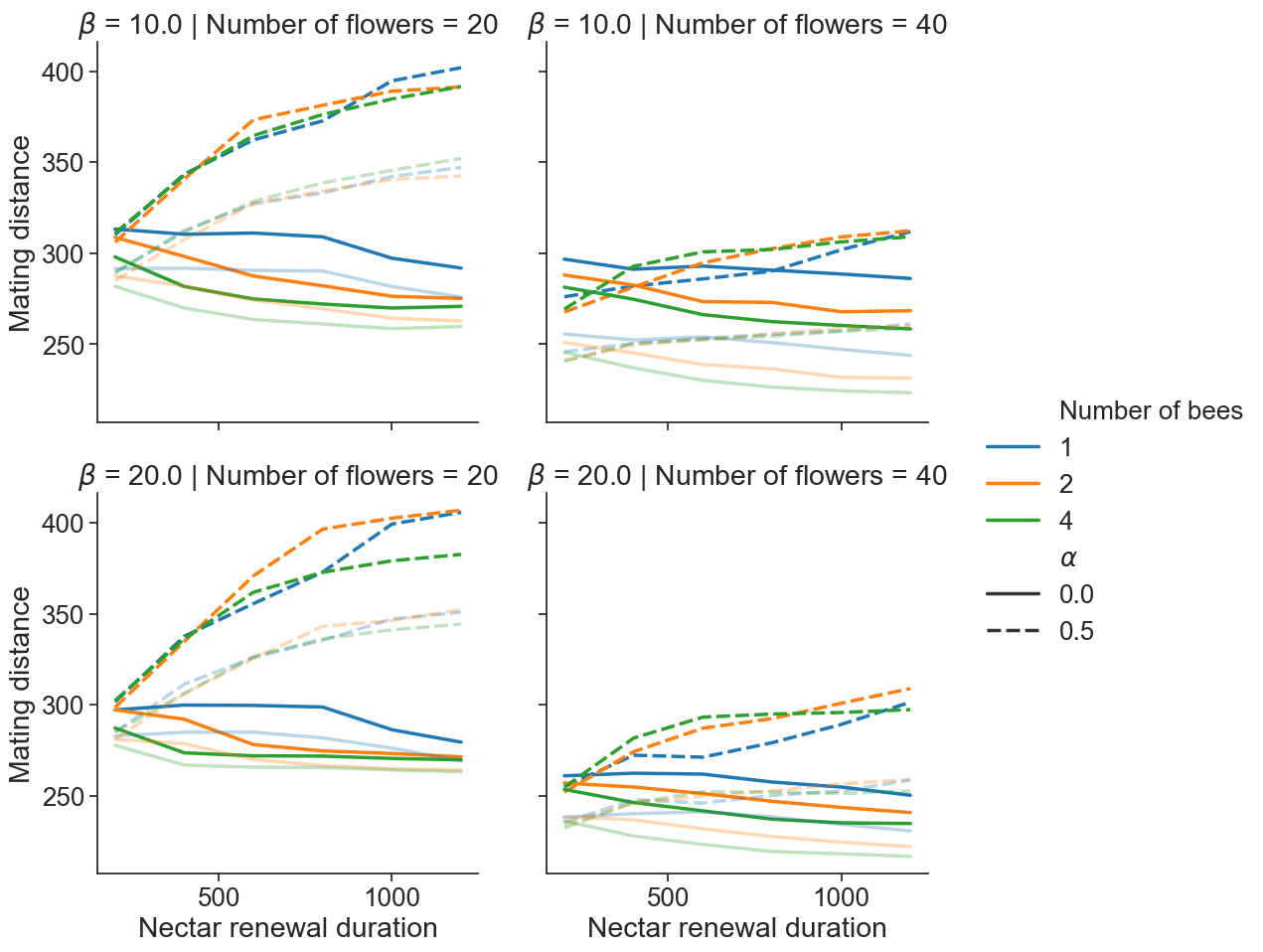

*Figure S5. Effect of learning rate* $\alpha$ *(line style), exploration-exploitation parameter* $\beta$ *(row), number of flowers (column), number of bees (line color), pollen deposition rate (*$p=0.25$ *for opaque lines and* $p=0.5$ *for transparent lines), and nectar renewal duration (x-axis) on median plant population mating distance (y-axis).*

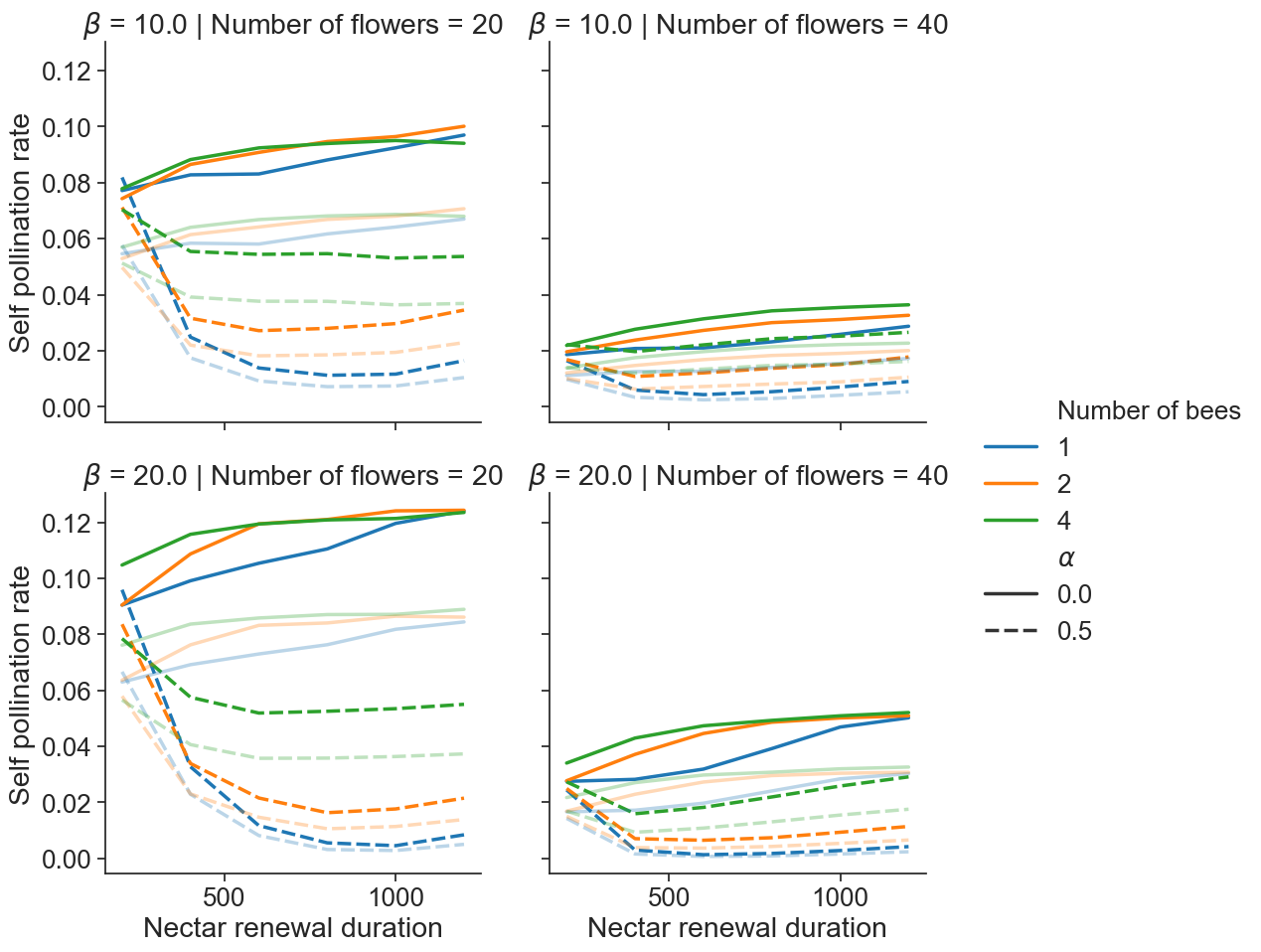

*Figure S6. Effect of learning rate* $\alpha$ *(line style), exploration-exploitation parameter* $\beta$ *(row), number of flowers (column), number of bees (line color), pollen deposition rate (*$p=0.25$ *for opaque lines and* $p=0.5$ *for transparent lines), and nectar renewal duration (x-axis) on average self-pollination rate of a plant (y-axis).*

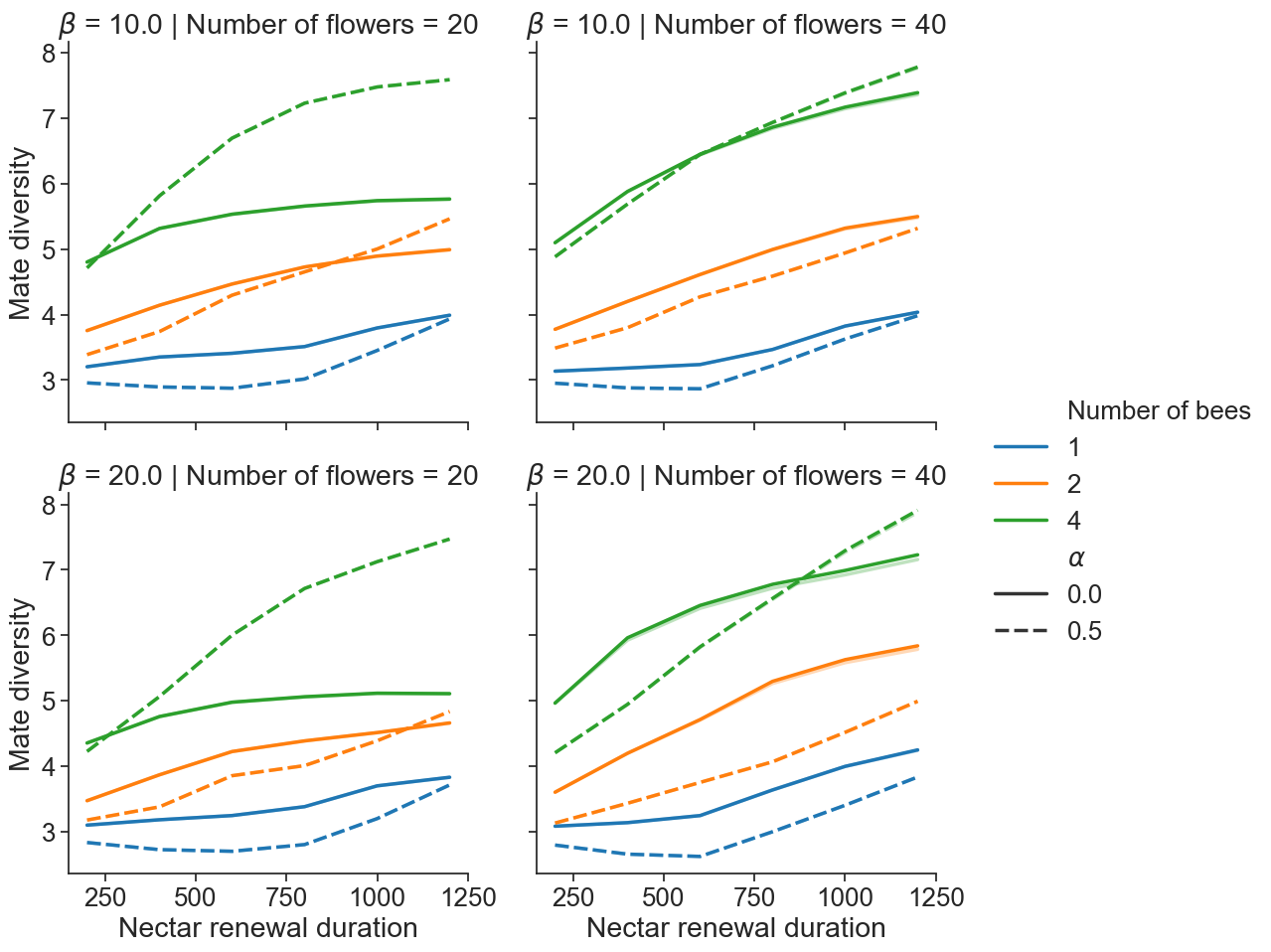

*Figure S7. Effect of learning rate* $\alpha$ *(line style), exploration-exploitation parameter* $\beta$ *(row), number of flowers (column), number of bees (line color), pollen deposition rate (*$p=0.25$ *for opaque lines and* $p=0.5$ *for transparent lines), and nectar renewal duration (x-axis) on average mate diversity of a plant (y-axis). The pollen deposition rate did not affect mate diversity values which is why the transparent lines are not visible on the graph.*

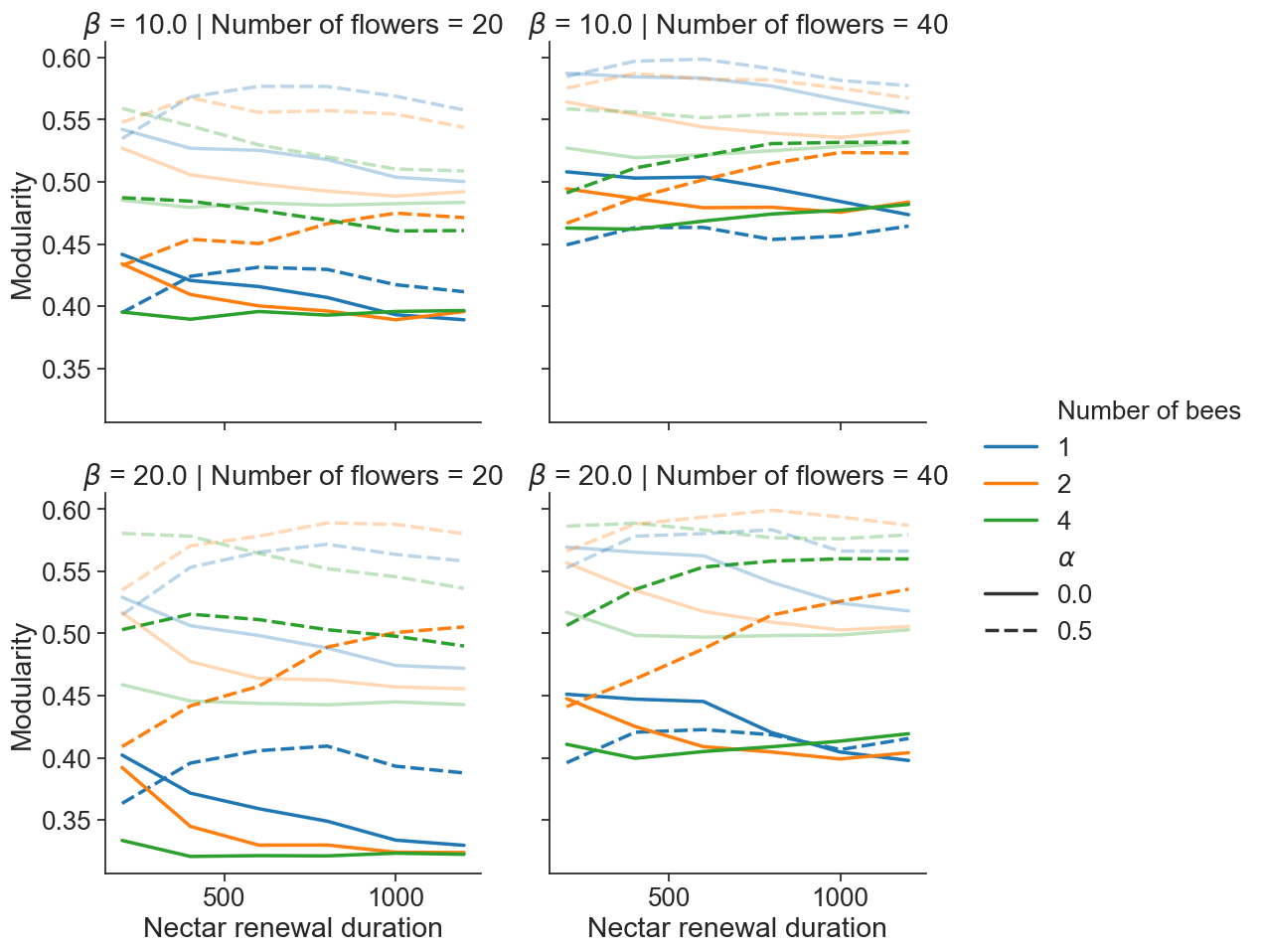

*Figure S8. Effect of learning rate* $\alpha$ *(line style), exploration-exploitation parameter* $\beta$ *(row), number of flowers (column), number of bees (line color), pollen deposition rate (*$p=0.25$ *for opaque lines and* $p=0.5$ *for transparent lines), and nectar renewal duration (x-axis) on average modularity of the donor-recipient plant bipartite network (y-axis).*
